# Designing function-specific minimal microbiomes from large microbial communities

**DOI:** 10.1101/2023.06.12.544531

**Authors:** Aswathy K. Raghu, Indumathi Palanikumar, Karthik Raman

**Affiliations:** Centre for Integrative Biology and Systems mEdicine (IBSE), Indian Institute of Technology (IIT) Madras, Chennai — 600 036, India; Robert Bosch Centre for Data Science and Artificial Intelligence (RBCDSAI), IIT Madras, Chennai 600 036, India; Department of Biotechnology, Bhupat Jyoti Mehta School of Biosciences, IIT Madras, Chennai 600 036, India

## Abstract

**Motivation:** Microorganisms thrive in large communities of diverse species, exhibiting various functionalities. The mammalian gut microbiome, for instance, has the functionality of digesting dietary fibre and producing different short-chain fatty acids. Not all microbes present in a community contribute to a given functionality; it is possible to find a *minimal* microbiome, which is a subset of the large microbiome, that is capable of performing the functionality while maintaining other community properties such as growth rate. Such a minimal microbiome will also contain keystone species for SCFA production in that community. In the wake of perturbations of the gut microbiome that result in disease conditions, cultivated minimal microbiomes can be administered to restore lost functionalities.

**Results:** In this work, we present a systematic algorithm to design a minimal microbiome from a large community for a user-proposed function. We employ a top-down approach with sequential deletion followed by solving a mixed-integer linear programming problem with the objective of minimising the *L*_1_-norm of the membership vector. We demonstrate the utility of our algorithm by identifying the minimal microbiomes corresponding to model communities of the gut, and discuss their validity based on the presence of the keystone species in the community. Our approach is generic and finds application in studying a variety of microbial communities.

**Availability:** The algorithm is available from https://github.com/RamanLab/minMicrobiome

**Author Summary:** Microorganisms are ubiquitous in nature. They survive in communities by interacting with each other and influence the biosphere by carrying out specific functions. For instance, the mammalian digestive system is heavily dependent on microbial communities in the gut (known as gut microbiome) to digest dietary fibres which are otherwise indigestible. The capability of gut microbes to convert dietary fibres to short-chain fatty acids help the host by regulating the functionality of the gut epithelial barrier. Oftentimes, some members of a community have redundant functions. Hence, it is possible to find a smaller subset of organisms that is capable of a given functionality, while also maintaining the required growth rate. We call them a minimal microbiome. Knowledge of such function-specific minimal microbiomes is useful for constructing communities for laboratory study and for designing treatment strategies for medical conditions caused by microbiome disruption. We present an optimization algorithm for identifying function-specific minimal microbiomes from a large community. We also demonstrate the performance of the algorithm by analysing minimal microbiomes obtained from some known communities. Overall, our research work highlights the significance of function-specific minimal microbiomes and provides an efficient computational tool for their identification.

## Introduction

Microorganisms seldom exist solo in nature; they form communities and survive by interacting with other microbial species. Communities are known to exhibit cooperation and competition depending on the composition of member species and environmental conditions [1]. The human gut harbours at least 1000 different species, comprising 10^13^–10^14^ organisms, whose collective genome is at least 100 times the human genome [2]. The gut microbiome is an essential part of the digestive system that synthesizes amino acids and vitamins and breaks down the otherwise indigestible dietary fibre into products that the human body can absorb [3, 4]. It has a massive influence on human health, and its disruption has been found to cause several disease states [5] such as obesity [6, 7], type II diabetes [8], and inflammatory bowel disease [6]. Microbes in the gut are known to co-exist by cross-feeding metabolites [9] and by performing complementary metabolic functions [1].

Gut microbes predominantly survive on the dietary fibre, glycans and secretions from the host epithelial cells [10]. The symbiotic association between humans and the gut microbiome is demonstrated through the role of microbiome-derived short-chain fatty acids (SCFA), bile acids and other small molecules in maintaining energy homeostasis and regulating gut barrier and inflammation [11]. Especially, SCFA has involved in adaptive immune system response, energy for the growth of colon epithelial cells, cholesterol synthesis and crosstalk with other tissues like the lung and liver [3, 12, 13]. To give an example, butyrate has been reported to have a major influence on maintaining host health owing to its ability to induce apoptosis [14], develop intestinal barrier [15] and regulate the immune system [16]. Perturbation to the community, such as that brought about by antibiotic usage [17–19], can disturb the community and affect its functionality. All the members of the microbial community do not contribute equally to the production of all the metabolites. Only some of them have the capability to break down dietary fibre, and such species are regarded as keystone species in the community [20, 21]. For instance, certain species of *Bifidobacterium, Bacteroides* and *Firmicutes* have the ability to degrade polysaccharides [22]. The numerical abundance of such organisms may not be significant compared to their functional impact on the community [20, 21]. Microbiomes also possess functional redundancy due to which dissimilar organisms capable of similar functions can be interchanged [21]. Understanding the interaction characteristics of the member species is paramount to comprehend the features of the overall microbiome, and several works have been published in this regard [20, 23].

Computational tools are very effective and widely used for studying the behaviour of microbes in a community using their genome-scale metabolic models. Numerous approaches at varying levels of complexities are available for microbial community modelling [24–29]. Previous literature has shown that synthetic microbial communities comprising representative species from major phyla present in the gut [30] can be used for treating diseases such as *Clostridium difficile* infection (CDI) and Inflammatory Bowel Disease (IBD) [31]. Algorithms such as MiMiC [32] use data-driven approaches to define synthetic communities consisting of all the identifiable members, as a proxy for native communities. Constraint-based modelling methods are useful for quantitative predictions of community features [24–27, 29].

In this work, we present a systematic mathematical approach to design functionality-dependent minimal microbiomes of a given large community. The minimal microbiome so identified need not be unique—there can be many possible combinations of species that form a minimal microbiome. Knowledge of such minimal microbiomes would be useful in designing treatment strategies for diseases caused by a disruption in the gut microbiome. For example, instead of a faecal transplant to treat *Clostridium difficile* infection, a minimal microbiome with the required functional capability of a large healthy microbiome can be cultivated and administered. This would in turn reduce the risk of unintentional transfer of pathogenic microbes [33]. Microbial communities play an important role in bioengineering applications such as chemical production [34] and waste-water treatment [35, 36], and the concept of the minimal microbiome to recover from perturbations is useful in those situations as well. This work identifies functionality-specific model microbiomes using a constraint-based approach, which could be considered as the ‘community’ analogue of the minimal reactome of an organism [37]. It helps to develop customized communities based on the application. Eng & Borenstein [38] have published a network-based approach to this problem, by identifying the pathways from substrates to products and minimizing the number of organisms needed for catalyzing the reactions using integer linear programming (ILP) formulation. Our algorithm is flexible and allows one to choose the desired functionality, such as the maximization of the production of a metabolite or the sum of a few metabolites.

## Results and Discussion

### Analysis of a 9-member community for butyrate production

A complex web of cross-feeding between the microbial species in a community is required to digest the dietary substrates that reach the colon undigested, and to produce SCFA. Despite their intricate exchanges, gut microbiome present high functional redundancy and environment-specific variation in inter-species interactions. Butyrate is a crucial SCFA that provides several health benefits to the host due to its anti-carcinogenic [14] and anti-inflammatory properties [16]. We analyzed a model community with the functionality constraint on maximum butyrate production (Constraint 3) for the gut microbiome consisting of 9 organisms [24] namely *Bacteroides thetaiotaomicron* VPI 5482 (*Bt*), *Eubacterium rectale* ATCC 33656 (*Er*), *Faecalibacterium prausnitzii* A2 165 (*Fp*), *Enterococcus faecalis* V583 (*Ef*), *Lactobacillus casei* ATCC 334 (*Lc*), *Streptococcus thermophilus* LMG 18311 (*St*), *Bifidobacterium adolescentis* ATCC 15703 (*Ba*), *Escherichia coli* SE11 (*Ec*) and *Klebsiella pneumoniae pneumoniae* MGH78578 (*Kp*), on AGORA high fibre and Western diets and identified their minimal microbiomes. The parameter values used are 0.8 for both gr_frac and scfa_frac and 0.99 for gr_opt_frac. The results of the simulation are tabulated in Table 1.

**Table 1.**
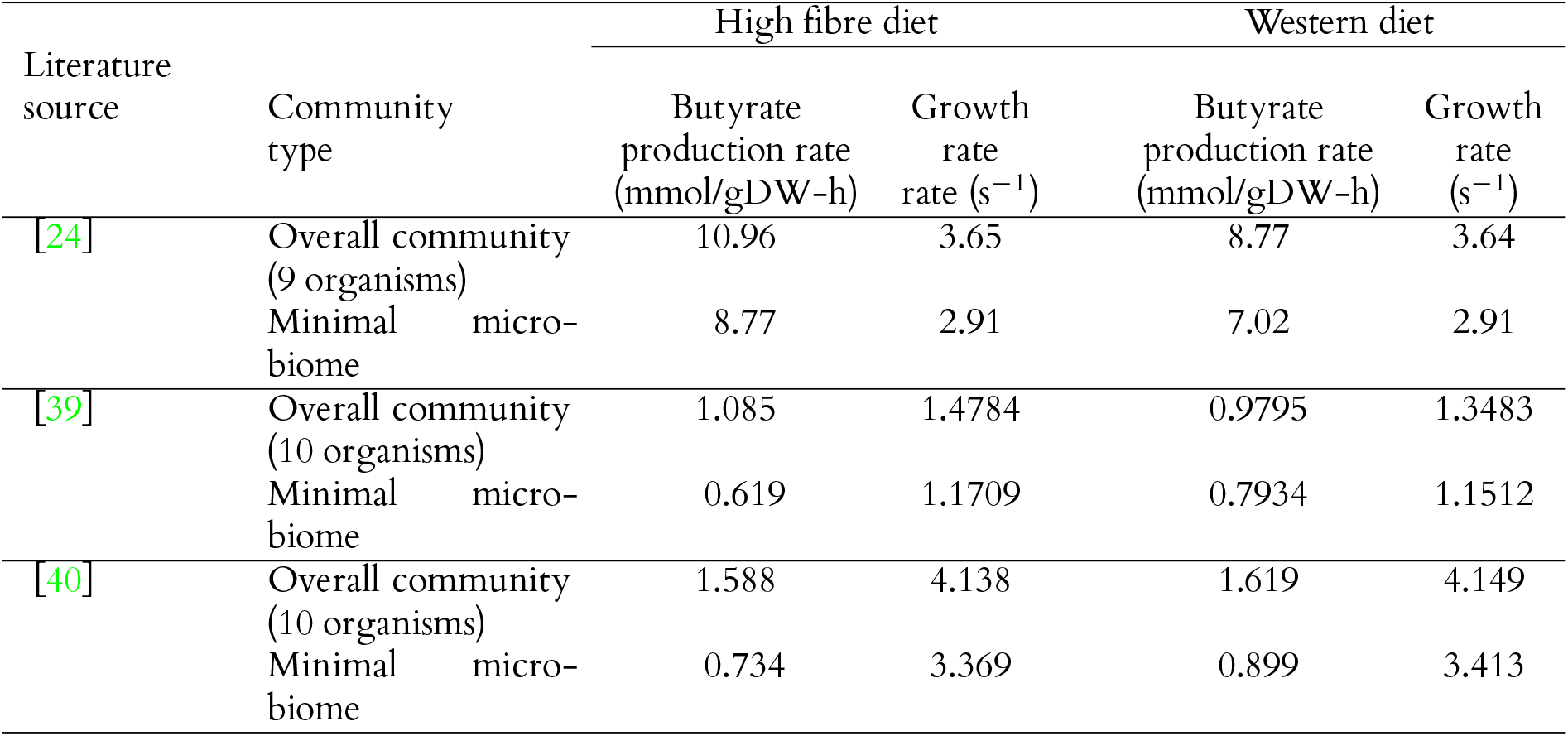
Simulation results based on [24], [39] and [40]

On a high-fibre diet, the overall growth rate of this community was calculated to be 3.65 h^−1^, and the individual growth rates were [0.29, 0.007, 0.006, 0.000856, 0.065, 0, 0.27, 2.98, 0.027] h^−1^. The maximum butyrate produced by this community when the growth rate of each species is constrained to be at 99% (gr_opt_frac) of the above individual growth rates was 10.96 mmol/gDW-h. The minimal microbiome that has at least 80% growth (gr_frac) and butyrate production rates (scfa_frac) as that of the above community (Constraint 3) consists of only *Fp* A2 165 and *Ec* SE11. This minimal microbiome can grow at the rate of 2.91 h^−1^ while producing butyrate at 8.77 mmol/gDW-h. Since sub-optimal growth rates increase SCFA production, the constraint of 80% on the growth rate makes the above solution possible. If the two organisms, *Fp* and *Ec*, were to be simulated as a community maximizing their growth rate, a higher growth rate with lower butyrate production would have resulted. Now, for calculating the maximum butyrate by the community, if 90% of individual growth rates (gr_opt_frac) is used instead of 99%, the butyrate production of the community is 18.79 mmol/gDW-h, with a minimal microbiome comprising *Bt, Fp* and *Kp*, that produces 15.03 mmol/gDW-h butyrate. On the other hand, if the lower bound for growth rate (gr_frac) is 90% instead of 80%, the minimal microbiome will have 3.28 h^−1^ growth rate with 8.77 mmol/gDW-h of butyrate production. For this case, there are multiple possible minimal microbiomes containing 3 organisms: *Bt, Ec* and *Kp* or *Er, Fp* and *Ec*. The same community (with parameters gr_frac = scfa_frac = 0.8 and gr_opt_frac = 0.99) on a Western diet, has a growth rate of 3.64 h^−1^ with a butyrate production rate of 8.77 mmol/gDW-h. The minimal microbiome is *Fp* and *Ec*, with a growth rate of 2.91 h^−1^. On the Western diet, butyrate production is comparatively lesser, yet the same organisms constitute the minimal microbiome, which is in accordance with the reduction of SCFA in high-fat diets [13].

It may be noted that all the minimal microbiomes identified consist of either *Fp* or *Bt*, which are the organisms capable of digesting dietary fibre, and are thus the keystone species, in the above community [24]. The presence of either of these species at different fibre uptake rates has been reported in literature [24] and hence the above findings support our algorithm’s predictions. Some species, such as the keystone species, could be interchangeable but not essentially equal [21]. The algorithm is not to find all the abundant species in the community, but to find the possible minimal community that includes the keystone species for a specific purpose. The keystone species need not be an abundant species in the microbiome.

The constraint on maximizing the sum of SCFAs (Constraint 1), although intended to find the maximum total SCFA production, sometimes results in the production of only acetate and propionate because the production of acetate is the highest and thus maximizes the objective. The code has been enabled for a weighted sum of SCFAs where weight can be provided by the user. The aforementioned community of 9 organisms was found not to produce acetate and butyrate simultaneously. If the weightage used in the sum of acetate, butyrate and propionate is 1:1:1 or 1:2:1, the production will be 24.5540, 0, 0.9063 mmol/gDW-h, respectively. On the contrary, if the weightage is 1:3:1, the corresponding fluxes are -1.0000, 10.9582, 0.9063 mmol/gDW-h. If acetate in the diet is removed (by changing the lower bound to 0 from -1), the corresponding fluxes when the weightage is 1:3:1 are 0, 10.7317, 0.9103 mmol/gDW-h, respectively. There is only a slight dip in the production of butyrate on the removal of acetate from the diet.

Not all the solutions to the minimization problem, but only those ones containing the least number of species, are referred to as minimal microbiomes. A minimal microbiome does not identify all the keystone species, so it may be possible to have other smaller microbiomes (if not minimal), capable of the same functions. Multiple minimal microbiomes are also possible because multiple combinations of the member species might satisfy the imposed functionality constraints. They could even be mutually exclusive depending on the type of member species. Thus, the deletion of one of the minimal microbiomes from the large community need not affect the growth rate and functionality of the latter.

### Identification of diet-based minimal gut microbiome

A recent study [39] identifies 10 microbial species as a Diet-based Minimal Microbiome (DbMM) for the effective conversion of dietary fibres to short-chain fatty acids (SCFA). The study employs an eco-physiology-guided [41] approach to identify SCFA-producing (butyrate, propionate) stable minimal microbiome that could utilise multiple dietary substrates. The identified minimal microbiome embodies functional modules involved in complex carbohydrate degradation to simple sugars and fermentation products and the production of SCFA using the degraded products. Minimal microbiomes and key species under a host-based diet from DbMM are evaluated to ensure the ability of the proposed algorithm.

Minimal microbiome from 10 core microbial species [*Faecalibaterium prausnitzii, Coprococcus catus, Bacteroides ovatus, Bacteroides xylanisolvens, Agathobacter rectalis, Anaerobutricum soehngenii, Eubacterium siraeum, Flavonifactor plautii, Roseburia intestinalis*, and *Subdoligranulum variable*] is investigated under different host-based diet conditions (western and high fibre diet) for maximum SCFA (butyrate and propionate) production with optimal community growth. A 10-member microbial community under the western diet exhibits a growth of 1.35 h^−1^ with 0.921 mmol/gDW-h of propionate production. Butyrate is not produced by the community while optimised for biomass. Maximum butyrate production of 0.979 mmol/gDW-h is attained when the growth rate of the community (gr_frac) and individual species (gr_opt_frac) in a community is constrained to 80% and 99%, respectively. A 2-member community of *Bacteroides ovatus* and *Coprococcus catus* is considered a minimal microbiome since they could produce more than 80% of the maximum SCFA produced by a 10-member community. Since the objective is to maximise the sum of different SCFA metabolites (propionate and butyrate), the solution obtained showed only propionate production. At a weighted sum of SCFA (propionate: butyrate - 1:1), 0.921 mmol/gDW-h of propionate production is observed with no butyrate production, and the butyrate production of 0.979 mmol/gDW-h at a ratio of 1:5. While optimising for butyrate production, the minimal microbial community includes *B. ovatus* and *C. catus* along with either *Fp* or *Eubacterium* sp. The community captures the cross-feeding of species capable of degrading complex carbohydrates to simple metabolites, such as lactate (*B. ovatus* [42, 43]) and a species which could convert the simple metabolites to propionate (*C. catus* [44]) or butyrate (*Fp* [45], *Er* [46]).

The reduced constraint for the growth of individual species (gr_opt_frac) in a community to 0.9 instead of 0.99 improves the maximum butyrate production (Constraint 2) to 9.436 mmol/gDW-h (≈ 10-fold compared to the results for gr_opt_frac = 0.99), and 4.606 mmol/gDW-h of butyrate production is detected at a ratio of 1:2 (propionate: butyrate). However, the growth of the community drops down to 1.215 h^−1^. Keystone species of the microbial community are consistent, and all three species (*B. ovatus, C. catus* and *E. rectale*) are identified as part of the minimal microbiome that facilitates maximum butyrate production.

The same pattern of butyrate and propionate production was observed when the community was simulated on a high-fibre diet. The core microbial community exhibited a growth of 1.479 h^−1^ with 0.901 mmol/gDW-h propionate. The maximum butyrate production of 1.085 mmol/gDW-h is observed at a ratio of 1:5 (propionate: butyrate). Though *B. ovatus, C. catus* and *F. prausnitzii* are the keystone species in a microbiome, *E. rectale* is replaced by *Subdoligranulum variabile* [47] as a fourth key species, a member of the minimal microbiome for butyrate production. A 10-fold increase in butyrate production (10.499 mmol/gDW-h) with decreased growth (1.346 h^−1^) is observed when the individual species’ growth in a community is constrained at 90%. The minimal microbiome analysis suggested that 10 microbial species can produce SCFA from various complex carbohydrates. The proposed algorithm could identify the minimum number of microbial species needed to produce SCFA (desired metabolites) for a specific carbon source.

### Minimal microbiome of a synthetic therapeutic consortium

A synthetic consortium is designed to treat inflammatory bowel disease (IBD) by complementing the missing critical functions of the human gut microbiome. A bottom-up approach-based rational consortium design identified a 17-species therapeutic consortium, including *Megamonas funiformis, Megamonas hypermegale, Acidaminococcus intestini, Bacteroides massiliensis, Bacteroides stercoris, Barmesiella intestinihominis, Fp, Subdoligranulum variabile, Anaerostipes caccae, Anaerostipes hadrus, Clostridium symbiosum, Akkermansia muciniphila, Clostridium scindens, Clostridium boltae, Blautia producta, Blautia hydrogenotropia* and *Marvinbryantia formatexigens* [40]. The consortium is analysed for its therapeutic function to support mucosal homeostasis (SCFA-mediated) and immune modulation defects (bile acid-mediated) in the gut microbiome modulated during the IBD condition. The synthetic community designed to treat IBD and promote gut health has high functional redundancy and complementary auxotrophies for better stability and efficacy. The minimal microbiome of the 17-species consortium for a specific objective will provide insights into the role of microbial species in a community, such as butyrate production, community resilience and secondary bile-acid production.

Here, the algorithm identifies key species involved in the SCFA (propionate and butyrate) mediated energy homeostasis in the microbiome [48]. Default growth parameters are applied to determine the minimum number of microbial species needed to produce the maximum SCFA from the community (gr_opt_frac = 0.99; gr_frac = 0.8; scfa_frac = 0.8). A synthetic consortium exhibits maximum butyrate production of 1.588 mmol/gDW-h and propionate production of 2.382 mmol/gDW-h at a growth rate of 4.138 h^−1^ on a high-fibre diet. Minimal microbiome reveals *C. boltae, M. funiformis* and *C. symbiosum* as key microbial species for SCFA production and can produce nearly 50% of the maximum butyrate (0.734 mmol/gDW-h). The butyrate production increases 13-fold when the growth rate of the community is constrained at 90% instead of 99%. Since the minimal microbiome can vary based on the sequence of the microbes removal from a community, *B. massiliensis*, a propionate producer, replaces the commonly observed propionate producer *M. funiformis* in a few iterations to form a minimal microbiome.

The higher growth rate of the butyrate-producing microbe, *C. symbiosum* [48] and the propionate-producing microbe, *M. funiformis* [49], is observed in the minimal microbiome than in synthetic consortium when it is analysed for growth and SCFA production potential. The results suggest that higher butyrate production (1.588 mmol/gDW-h) supported by the key species in 17-species consortia is complemented by the growth of the other microbial species in a community.

A similar trend of butyrate production and growth profile of a community and minimal microbiome on a Western diet is observed (Table 1). The minimal microbiome consists of *C. boltae, M. funiformis* and *C. symbiosum*, produces 0.899 mmol/gDW-h of butyrate, at a growth rate of 3.413 h^−1^ and 7.3278 mmol/gDW-h at a growth rate of 3.1028 h^−1^. Minimal microbiome identifies 3 key species to complement the SCFA production ability since the analysis focused on maximising SCFA with a constrained growth of the individual species (80%) and community (99%). Further analysis showed that the other microbial species support the community in fulfilling other significant functions, such as secondary bile acid production and better resilience to pathogen colonisation.

### Minimal microbiomes of synthetic communities

We now examine some of the known model microbial communities to identify their minimal microbiomes. [50] studied simplified human intestinal microbiota (SIHUMI) comprised of *Anaerostipes caccae, Bacteroides thetaiotaomicron, Bifidobacterium longum, Blautia producta, Clostridium ramosum, Escherichia coli* and *Lactobacillus plantarum*, complemented with *Clostridium butyricum* for butyrate production. They observed improvement in butyrate production with the modified SIHUMIx. The minimal microbiome for this community on a high-fibre diet, with the functionality to maximize butyrate, was found to consist of *Escherichia coli* and *Clostridium butyricum*. The butyrate production by the whole community is 15.16 mmol/gDW-h, whereas that by the minimal microbiome is 12.13 mmol/gDW-h. *Clostridium butyricum* is an irreplaceable member of the minimal microbiome as deletion of this species from the community reduces butyrate production to 11 mmol/gDW-h, which violates the constraint of at least 80% butyrate production.

A community consisting of 25 organisms with a high-fibre diet was studied, and 18 minimal microbiomes were identified for butyrate production. Fig 1 shows the frequency at which a given species in the microbiome was identified as a part of a minimal microbiome. Some species of *Bacteroides* are capable of breaking down glycans [22], and thus show up very frequently as a part of the minimal microbiome. Due to the functional redundancy offered by several species, several minimal microbiomes with distinct species are possible and are indicated by the lower frequency appearances. A table showing a large community of 50 organisms and one of its minimal microbiomes on a Western diet with a constraint on butyrate production is given in Supplementary File S2.

**Fig 1.**
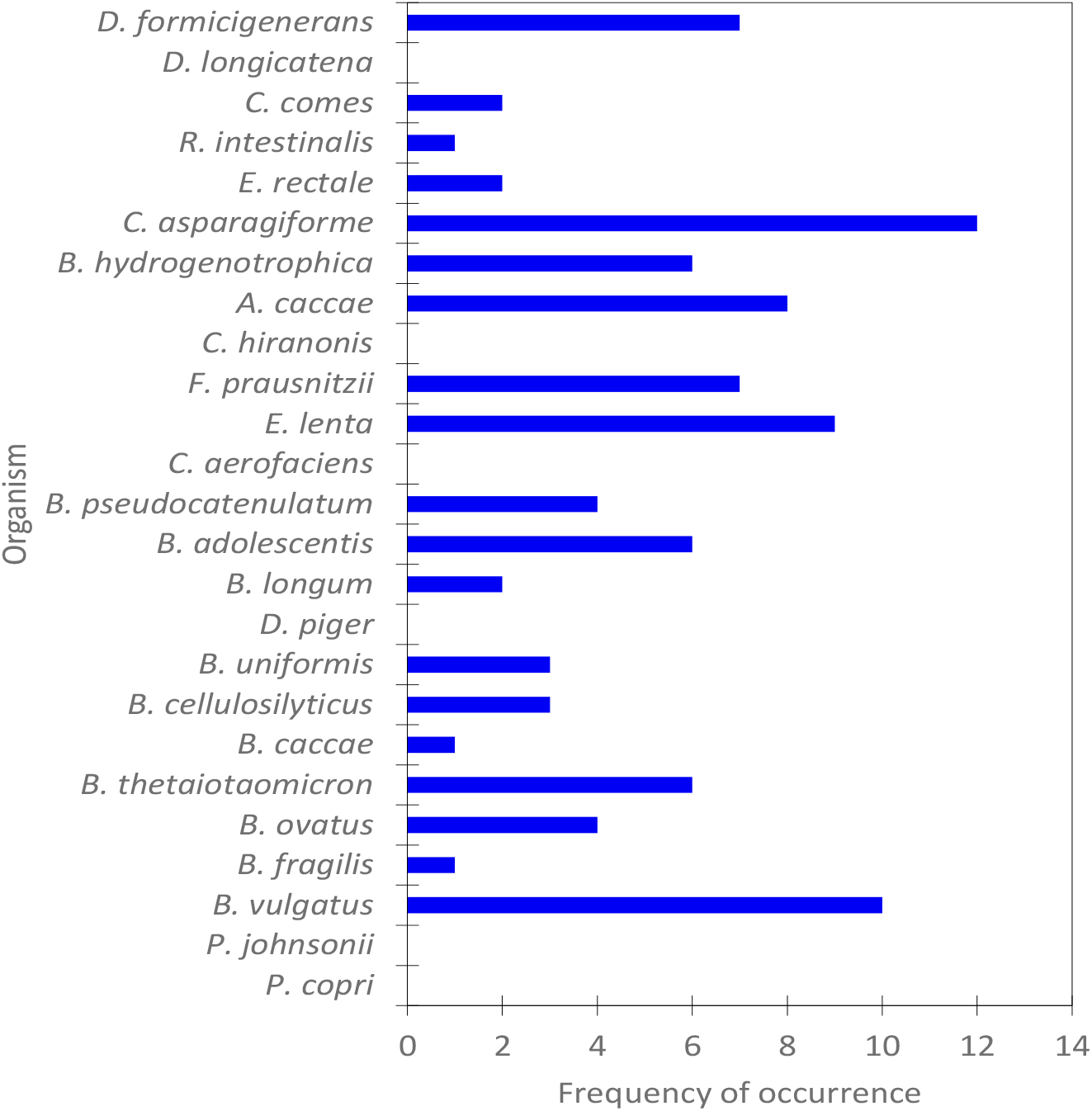
Frequency of occurrences of species in minimal microbiomes of a 25-member community on a high fibre diet. 18 minimal microbiomes are indicated.

### Performance of the algorithm

Running the MILP alone for a large community would be computationally intensive. However, the deletion of one species involves running an LP twice—one for checking the growth rate (FBA) and another (FVA) for the SCFA(s) produced. We observed that when smaller communities were considered, computational times were comparable when solved as an MILP or when size-reduction was done before MILP. For instance, when a 15-member community was run as an MILP (with 15 integer variables and the corresponding continuous flux variables), the time taken for computation was 101 s, whereas when it was reduced to 8 member community before running MILP, the time taken was 106 s owing to the additional number of LPs. The computations were done on a 2.40GHz 11th Gen Intel i5-1135G7 processor with 16 GB DDR4 RAM running Windows 11. However, for large communities, the size-reduction resulted in significant savings of computational time. MILP of a 25-member community took 1497 s while reducing it to 15 and 8 members reduced the time required to 430 s and 251 s, respectively, on the same machine. It must be noted, however, that the computational time also depends on the complexity of species models.

A known drawback of the algorithm is that it forces the metabolite cross-feeding between the organisms to attain the maximization objective; nonetheless, it is an admissible assumption in the case of the gut microbiome since they are known to be a co-occurring, cooperative community. If the spatial arrangement of the species or/and the regulatory constraints are known, the individual uptakes for each species can be accordingly adjusted for a more realistic solution. Besides, sub-optimal growth rates are known to result in better SCFA production, and the parameters may be adjusted to suit realistic conditions. In short, the code is flexible to account for a variety of scenarios. The algorithm, however, does not exhaustively identify all the possible minimal microbiomes, especially when multiple functional redundancies are present in the community. While it is readily possible to achieve that by using integer cuts to the MILP problem, there will be a definite drop in performance.

## Conclusion

Given a large microbial community, all of its members would not be contributing to certain specific functionalities of the overall microbiome. It is possible to design ‘minimal microbiomes’ specific to certain functionalities, and we present a simple, customisable constraint-based approach for identifying them. Such minimal microbiomes would contain keystone species of the community. Multiple minimal microbiomes, which may even be mutually exclusive, can exist with the capability for a given functionality. Knowledge of minimal microbiomes will be useful in designing microbial composition to treat certain diseases caused by disruption of the gut microbiome. The usage of a rationally designed community for treatment is better than faecal microbial transplantation from a donor that poses a risk for accidental pathogenic infection. The idea of a minimal microbiome can also be made use of in bioengineering applications involving microbial communities, to rescue them from undesired consequences of perturbations.

We presented a procedure of sequential deletion followed by solving a mixed integer linear programming problem (MILP) for the identification of a minimal microbiome having specific characteristics of the large microbiome considered. Being a constraint-based approach, the algorithm can quantitatively predict fluxes and metabolic exchanges and is reasonable for a co-existing community such as the gut microbiome. The algorithm has been designed to be flexible with the possibility of incorporating user-defined inputs for most constraints. We used this algorithm to identify the minimal microbiomes and keystone species in some model communities of the gut. The identified minimal microbiomes are found to be rational, and includes species that digest dietary fibre.

## Methods

### Formulation

Flux Balance Analysis (FBA) [51, 52] is an effective method for finding steady-state flux distributions in organisms using their genome-scale metabolic models. Joint FBA [53] is an extension of FBA for modelling microbial communities by a compartmentalized approach. It solves a linear programming (LP) problem with the defined objective, stoichiometric balances as the equality constraints and lower and upper bounds on fluxes as the inequality constraints. The formulation of joint FBA is as follows:

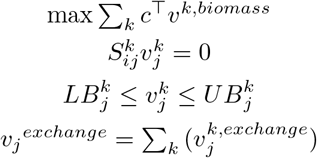

where *k* denotes species, *j* denotes reactions, *i* denotes metabolites, *S* is the stoichiometric matrix, and *v* are the reaction fluxes. In this approach, the overall exchange of each metabolite by the community is the sum of the corresponding exchange reactions of each organism, and these constraints enable metabolite uptakes and cross-feeding. However, in this approach, if the objective is to maximize the sum of biomass reaction fluxes of each organism, it could result in metabolite production without growth for certain organisms, which is unrealistic. This problem is surmounted by coupling the metabolic reactions of an organism to its biomass equation:

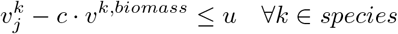

where *c* is the coupling vector and *u* is the threshold.

Due to the mathematical formulation to maximize the sum of biomass, constraint-based approaches such as joint FBA might force metabolite cross-feeding to attain the maximum possible growth. For co-occurring species in the gut microbiome, this cross-feeding would be reasonable, unless the species are known to be regulated differently or spatially separated.

To find the minimal microbiome, we define a binary membership vector identifying the status of membership of each species as 0: absent, or 1: present. This integer vector is used in the biomass constraint. The problem is now a mixed integer linear programming problem (MILP) and is solved with the objective of minimizing the *L*_1_-norm of the membership vector with the required functionality constraints. The following approach is used to find the functionality constraints:

1. The overall and individual growth rates of the given large community are calculated by solving the joint FBA LP problem.
2. The maximum possible rates of production of the desired metabolites—by default, SCFAs (acetate, butyrate and propionate), or their sum, is calculated by Flux Variability Analysis (FVA) by keeping the individual growth rates calculated in the previous step as the lower bounds for the individual biomass fluxes. The objective function for FVA is:

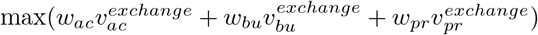

where *w* denotes the weightage for each of the fluxes. The default set of weightage is (1,1,1) and can be varied by the user if the production of one product is preferred to the other. The additional constraint on individual growth rates is given by:

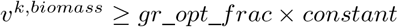

where the constant value is obtained from the joint FBA solution of step 1. If needed, a different metabolite list can be provided by the user in the optional inputs. The fraction of growth rates to be considered as lower bounds in this step can also be provided by the user (default value of ‘gr_opt_frac’ is 0.99). This value is relevant because a sub-optimal growth rate is known to be more realistic and results in good SCFA production [54]. In the code, the constraint on the sum of SCFAs is denoted as Constraint 1. Likewise, constraints on the production of acetate, butyrate and propionate are denoted as constraints 2, 3 and 4, respectively.
3. Fractions of individual growth rate calculated in Step 1 and SCFA production rate calculated in Step 2 are provided as the lower bounds for the corresponding fluxes in the MILP problem. The growth rate fraction (gr_frac) and SCFA production fraction (scfa_frac) can be provided by the user.

The MILP formulation is as follows:

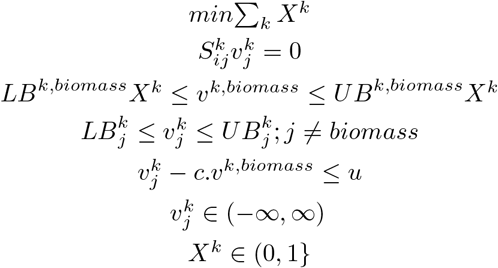

Functionality and growth rate constraints are captured as follows:

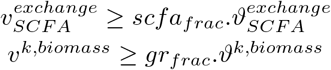

where *k* denotes species, *j* denotes reactions, *i* denotes metabolites, *S* is the stoichiometric matrix, *X* are the integer (binary membership) variables and *v* are the reaction fluxes (continuous variables). *LB* and *UB* denote the lower and upper bounds. *ϑ* in functionality constraints denotes reaction fluxes of the full community.

For large communities, solving an MILP is computationally challenging. In such cases, organisms are deleted one by one in a given sequence such that the functionality constraints are satisfied by the resultant community, until the community is small enough for reasonable computation for MILP solution. If the deletion sequence is not provided by the user, a random sequence is chosen by default. The deletion sequence is run over multiple times to reach the MILP size provided by the user. ‘Deletion’ of an organism is done by assigning 0 values to the lower and upper bounds of all the corresponding fluxes of the organism. However, the resultant minimal microbiome will depend on the sequence used for deletion and hence multiple iterations of the algorithm help to identify different possible solutions. The number of organisms in the minimal microbiomes identified in different iterations may vary.

The algorithm is explained in step-by-step detail in the Supplementary File S1, and a schematic is provided in Fig 2. COBRA toolbox [55] functions are used in MATLAB 2020b for the computations. The exchange and biomass reactions are identified in the code by the names used in AGORA [56] models, and each individual organism in the community is identified as ‘_org<number>‘ as in the communities created using the COBRA function createCommModel(). Gurobi (version 9.1.2, Gurobi Optimization LLC, USA) is the solver used for solving both LP and MILP.

**Fig 2.**
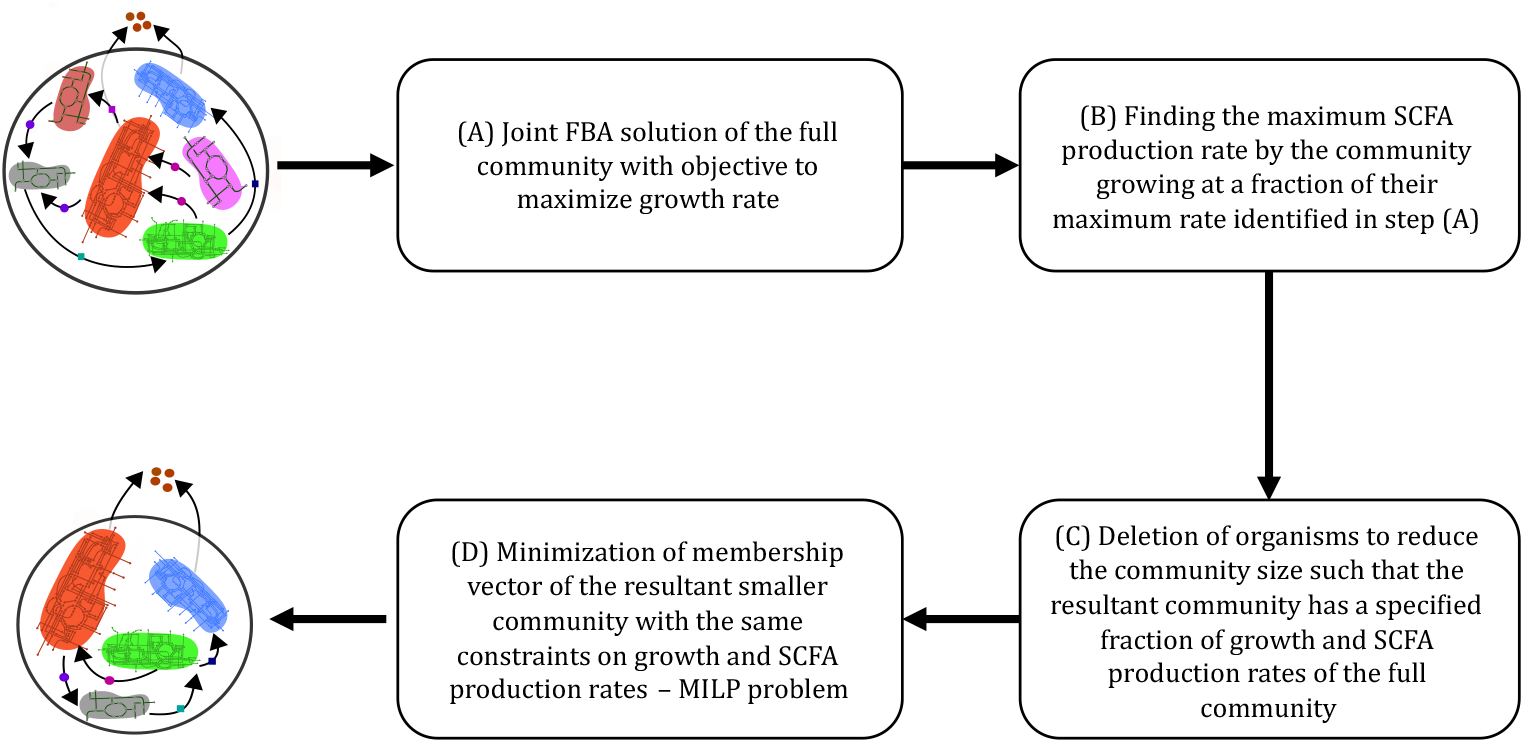
An overall schematic of the procedure followed to identify the minimal microbiome. Minimization is done in steps C and D, based on the constraints obtained from steps A and B.

## Acknowledgments

The authors thank L Raajaraam for useful comments on the manuscript.

## Financial support

AKR acknowledges support from the IBSE post-doctoral fellowship. IP acknowledges the Prime Minister’s Research Fellowship from the Government of India. KR acknowledges support from the Science and Engineering Board (SERB) MATRICS Grant MTR/2020/000490.

## Conflict of Interest

None declared.

## Notes

### Competing Interest Statement

The authors have declared no competing interest.

